# CHARACTERISATION OF FUNGI OF STORED COMMON BEAN CULTIVARS GROWN IN MENOUA DIVISION, CAMEROON

**DOI:** 10.1101/2020.10.31.363184

**Authors:** Teh Exodus Akwa, John M Maingi, Jonah K. Birgen

## Abstract

Common bean is a legume grown globally especially in developing countries including Cameroon for human consumption. In Cameroon it is grown in a wide variety of agro ecological zones in quantities enough to last through the off growing season after harvest. Stored common bean after harvest in Cameroon are prone to fungal spoilage which contributes to post harvest losses. This study aimed at characterising storage fungi on cultivars of stored common bean using morphological and molecular techniques. Fungi were isolated from six stored cultivars of common bean; Kidney bean, Black bean, Navy bean, Pinto bean, Pea bean and Large seeded bean plated on potato dextrose agar media. Cultural, micro-morphological and molecular techniques were used to characterise the isolates. A total of four fungi isolates resulted. Homology matches of the gene sequences in the Genbank databases identified the isolates to be; *Xylaria hypoxylon, Aspergillus flavus, Penicillium aethiopicum* and *Fusarium oxysporium*. Phylogenetic analysis and multiple sequence alignment showed these isolates were of distinct species. The species of fungi recovered from the stored cultivars signified poor preservation methods carried out after harvest. Effective management and control of these fungal species in common bean at storage will help reduce post-harvest losses and increase seed health.

## INTRODUCTION

Common bean is a legume grown globally especially in developing countries including Cameroon for human consumption. It is the most widely grown staple food crop in Africa (Beebe et al 2013). It is one of the most important staple food for the majority of the Cameroon population (Kamtchoum et al 2018). This is probably because it is suited to wide variety of agro ecological zones and can be grown in quantities enough to last through the off growing season after harvest. It is used to make bean cake which is a staple food consumed by nearly all the communities in Cameroon. In addition to being a source of food, it is a source of income to farmers.

Post-harvest activities play a vital part in the security of farm produce. These activities comprises handling procedures after harvest, transportation, storage and up to the point of enabling the availability of the produce to end consumers. In Cameroon, when common bean is harvested it is stored in varied structures by different communities. Such structures include; wooden granaries, grass lined granaries and silos. The storage of common bean in Cameroon usually last for an average period of 3 months after harvest which is later made available to consumers in markets.

Post-harvest losses of stored common bean have been reported in the West Region of Cameroon (Shende and Lifeter 2017). Post-harvest loss of stored common bean is partly due to storage fungi (Richard 2007). Storage fungi in common bean has led to its spoilage, decrease in quality and also resulted to a decrease in the germination of the common bean seeds (Kator et al 2016). Consumption of poor quality beans can also bring about health impacts (Kumar and Kalita 2017). Some fungal species on stored common bean can also release mycotoxins which causes human mycotoxicoses upon consumption. Typical example of mycotoxic fungus specie found on stored bean include is *Fusarium* which produces fumonisin that causes oesophageal cancer in humans and also interferes with sphingolipid metabolism. Another example is *Aspergilus* which produces aflatoxins known to cause liver cancer (Omotayo et al 2019).

Hence an urgent need is required to identify the fungus on stored bean so as to develop effective control and management measures of beans in stores. Thus, the aim of the research undergone was to isolate and characterize storage fungi associated with common bean after three months of storage.

## MATERIALS AND METHODS

### Study site

The study took place in the Menoua Division, West Region of Cameroon. Cameroon is located in the central part of Africa. Cameroon is situated at latitude 4° 8ʺ 40N - 4° 11ʺ North of the equator and between longitude 9° 16ʺ 90E – 9° 17ʺ 45E of the Greenwich meridian (Bechem and Afanga 2017).

### Sample collection

A multi stage sampling was used in the study. At the first stage a purposive sampling technique was used to select 6 zones/ sub divisions in the Menoua division where common bean was cultivated, consumed and equally stored several months after harvest. This selection was done based on geographical evidence of common bean farming. The sampled subdivisions were; Dschang, Santchou, Nkong ni, Fokoue, Penka Michel and Fongo Tongo. The second stage involved random sampling of the common bean cultivating farmers in each of the sampled subdivisions. At the last stage convenience sampling technique was carried out in collecting the common bean cultivars from the sampled farmers’ stores.

Stored common bean cultivars were collected from farmers’ storage structures (gunny bags). Sampling was carried out by picking the common bean grains three times (top, middle and bottom) from their sampled respective cultivar storage bags using a 15 cm diameter polyvinyl chloride (PVC) bowl.

A total of 500 grams of each common bean cultivar formed the bulk sample from each subdivision. These bulk samples served as representative samples of the common bean cultivars for evaluation throughout the research. Six different types of common bean cultivars (Figure1) were collected namely; Kidney bean (small size red), Black turtle bean (black bean), Navy bean (white bean), Pinto bean (mottled brown bean), Pea bean (mottled red bean) and Large seeded bean (Large red).

**Figure 1:**
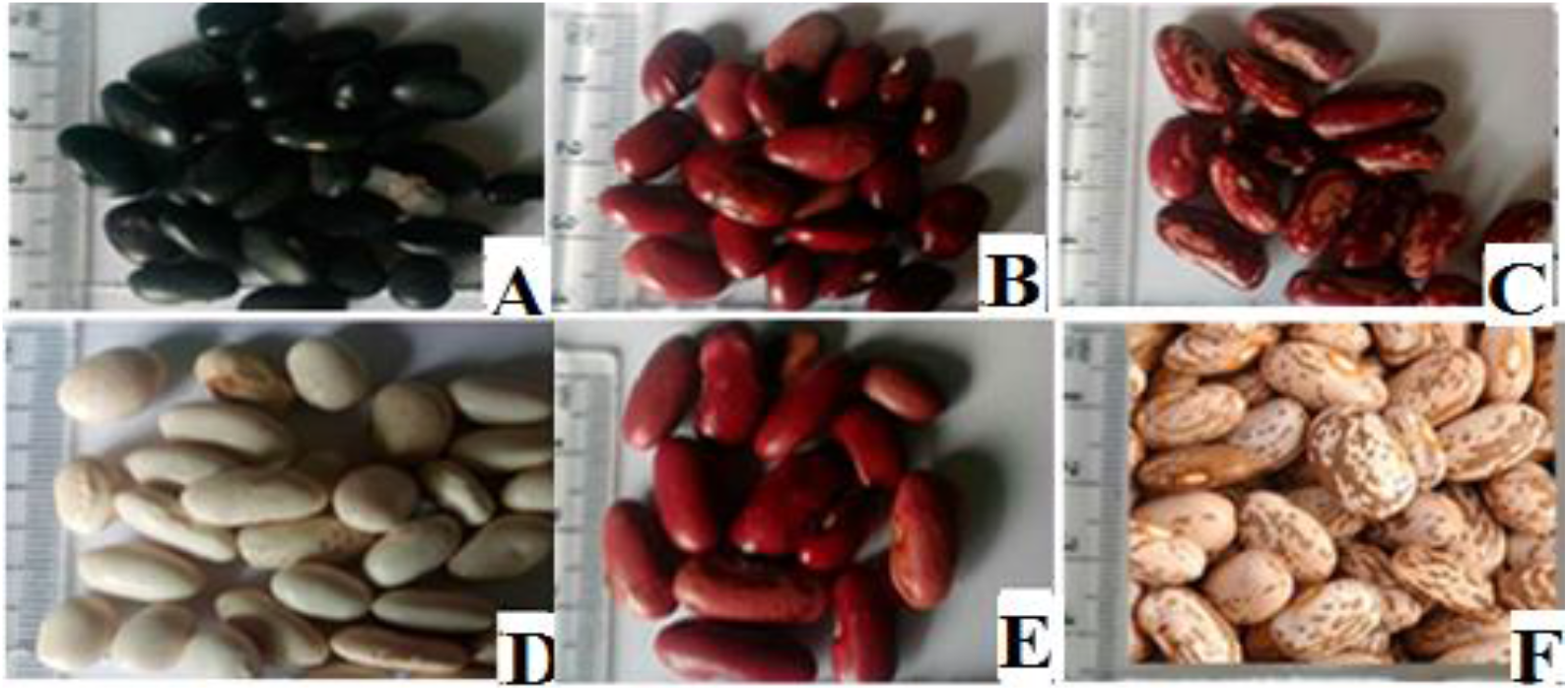
Cultivars of common bean collected from Menoua Division, Cameroon. A - Black turtle bean (Black bean) B- Kidney bean (small sized red bean) C- Pea bean (mottled red bean) D- Navy bean (White bean) E- Large seeded bean (Large red bean) F- Pinto bean (mottled brown bean)

### Detection and Isolation of Fungi from stored common bean cultivars

#### Surface sterilization of stored common bean cultivars

The common bean cultivars collected from the subdivisions were brought to the University of Buea Plant Science Laboratory, Cameroon for fungal isolation. A sub sample of 50 common bean grains was randomly taken from each bulk cultivar sample. These sub samples were surfaced sterilized for the maximum recovery of fungi using specific sterilization protocol (Schulz *et al* 1993). The common bean grains were immersed in 70 % ethanol for 1 minute. Surface sterilization of the grains was done using 10 % sodium hypochlorite solution for 1 minute. Finally, the grains were rinsed with sterile distilled water for 3 minutes. The common bean grains were then blot-dried with sterile filter paper.

#### Direct plating of bean cultivars for detection and isolation of fungi

The sterilised common bean grains obtained from the different cultivars were then plated on potato dextrose agar media using the agar plate method (Nega 2014). The plating of the common bean grains from each cultivar was done at a rate of 10 bean grains per plate (Figure 2) and was replicated five times. The grains were spread evenly in the plates. The plates were covered by their lids and fastened using parafilm. The sealed plates were then maintained under incubation at 28±2 °C for 7 days to promote fungal growth.

**Figure 2:**
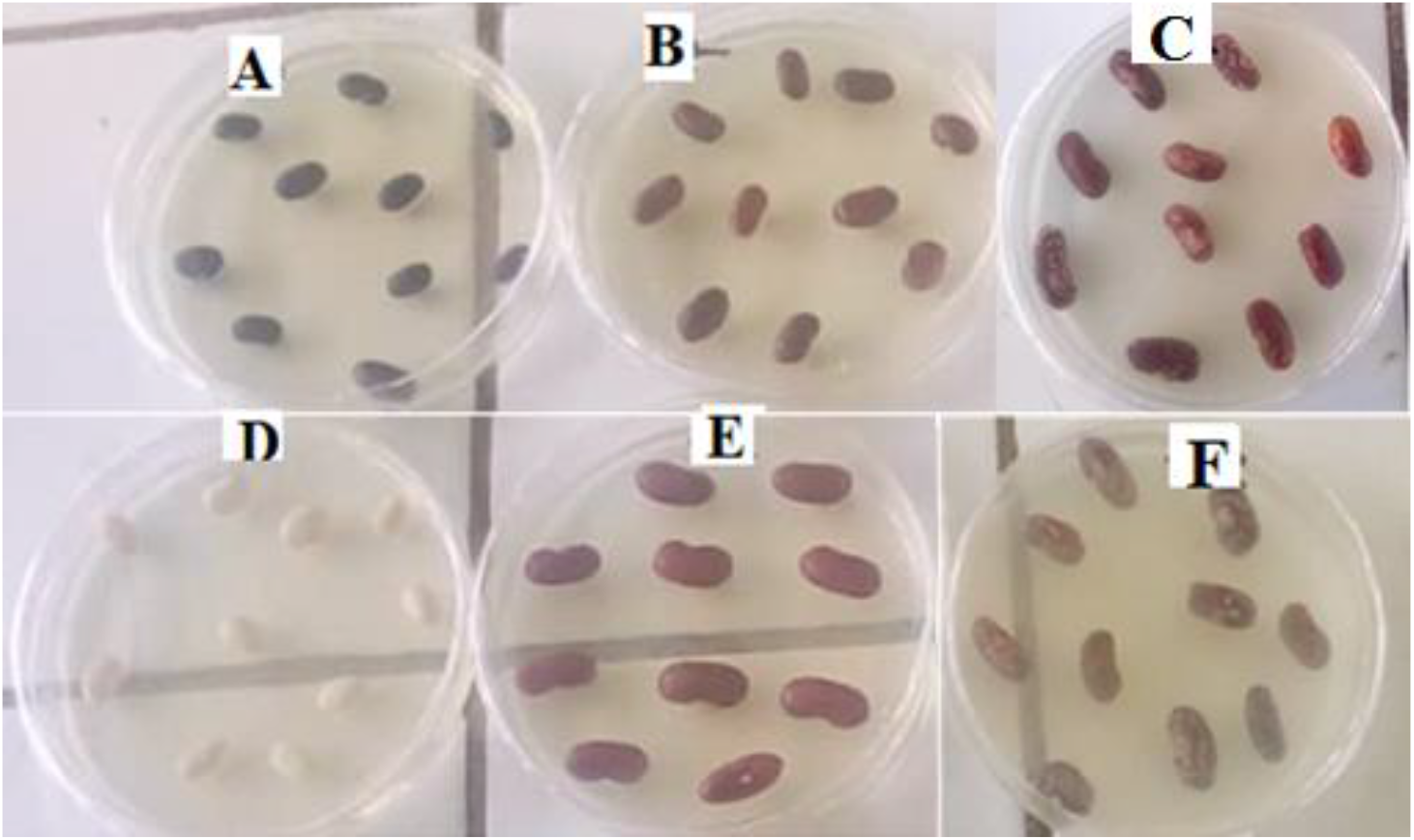
Common bean cultivars directly plated on PDA for isolation of fungi. A- Black turtle bean (Black bean) B- Kidney bean (small sized red bean) C- Pea bean (mottled red bean) D- Navy bean (White bean) E- Large seeded bean (Large red bean) F- Pinto bean (mottled brown bean)

#### Morphological Identification of Fungi from stored common bean cultivars

After 7 days of incubation, the plated common bean grains from each cultivar were observed for growth of fungi. Initial identification of fungal isolates was done using colony morphology and physical features according to Klich (2002). A loop of fungal mycelium was transferred with the use of a sterile inoculating needle from the bean grain showing fungal growth onto a fresh PDA culture medium in a petri dish. This was sealed then incubated at 28±2 °C for 7 days to obtain pure cultures.

For microscopic observation, a loop of fungal mycelium collected from the different fungal colonies with the use of a sterilized inoculating needle was placed on clean microscopic slides. A drop of lactophenol blue was added on each slide. A cover slip was gently placed on the slides. The excess lactophenol on the slides was blotted out. The slide were then mounted on a microscope stage and visualized through the x10 and x40 objective of the lens for the presence of sporulation and reproductive structures. Fungal cultures that could not sporulate were classified as ‘*mycelia sterilia*’ and sorted to morphospecies based on cultural characteristics (Lacap *et al* 2003).

Grouping of the fungal isolates was done based on similarities in their morphological characteristics shown by their colonies. Later, one isolate was picked out of each group as a representative for further analysis.

### Molecular characterisation of fungi on stored common bean cultivars

#### Fungal DNA Extraction

The extraction of the DNA from fungi obtained from the stored common bean samples was carried out following a standard protocol as described by Zhang *et al* (2010). This was done in the Molecular and Biotechnology Laboratory of the University of Buea. Thirty mg of mycelium from each representative isolate was scraped directly from the surface of the agar culture. The mycelium was then ground to a fine powder in an Eppendorf tube in liquid nitrogen using a pre-cooled pestle (Eppendorf no. 0030120973). The ground mycelium was resuspended and lysed in 500 ml of lysis buffer (40mmol/l Tris-acetate, 20mmol/l sodium acetate, 1mmol/l EDTA, 1% w/v SDS pH7·8). The tube and its content were then kept at room temperature for 10 minutes. Potassium acetate of volume 150 μl was later added into the eppendorf tube and then vortexed briefly, thereafter it was centrifuged at >13,000 x g for 1 minute. Another 1.5 ml eppendorf tube was used to collect the supernatant that resulted. To the supernatant, RNase A (10mg/ml) was added to get rid of RNA and then incubated at 37°C for 15 min. Isopropyl alcohol of volume 0.5 ml was then added to it. This was then followed by a 2 minutes centrifugation of the tube and its contents at >13,000 x g. Thereafter the supernatant was disposed. A volume of 0.3 ml of 70 % ethanol was used to wash the resulting DNA pellet in the eppendorf tube. The tube containing the pellet was then made to spin at 10,000 rpm for 1 minute and the supernatant disposed. The DNA pellet was then air dried and placed in 0.05 ml of Tris-EDTA to dissolve.

The presence of DNA was tested under 1.5 % agarose by gel electrophoresis. The extracted DNA was then stored under a temperature of −20 °C to maintain its stability.

#### PCR amplification of the ITS region of fungi

Amplification of the extracted fungal genomic DNA was performed using standard protocol as outlined by Ferrer *et al* (2001). The internal transcribed spacer (ITS) region of the ribosomal gene was used for species identification. For PCR reactions, the universal primer pairs ITS1/ITS4 (5′-TCC GTA GGT GAA CCT GCG G-3′/5′-TCC GTA GGT GAA CCT GCG G-3′) were used (Innis, 2012). The final PCR solution for each sample consisted of 25 μl of 2 × PCR master mix, 1 μl of each 2 μM primer, 1 μl of DNA (final concentration of 10 μM), and constituted to a final volume of 50 μl with nuclease free water. Polymerase chain reaction was performed using an eppendorf 96-well Thermocycler (Eppendorf, USA) and cycling conditions set as: 34 cycles denaturation at 98 °C for 1 min, annealing at 58 °C for 45 secs, extension at 72 °C for 1 min 30 seconds, final extension at 72 °C for 5 minutes and held at 4 °C until samples were retrieved.

Electrophoresis using 1% agarose gels was used to analyse the reaction product. The resulting size of the amplified PCR products by the ITS-1/ITS-4 pairs of primers was determined from the DNA molecular weight marker (200 bp marker).

#### Molecular sequencing of amplified fungal DNA

Sequencing of amplified fungal DNA was done at Inqaba Lab in South Africa. The Sanger dideoxy sequencing technology was used. DNA amplification was first confirmed using agarose gel electrophoresis. This was done with the use of a DNA ladder or marker. Alkaline phosphatase was used to clean the PCR products. The sequence reaction mixture included the purified PCR products, a big dye terminator V3.1 cycle sequencing kit with 3μl of DNA, a template and 1μl of 2μM of the same primer as used in the amplification process. Bi-directional sequencing of both DNA strands was done using the same forward and reverse primers used for amplification.

The established sequences of the ITS region were then deposited in the GenBank under respective accession numbers. For species identification, a BLAST analysis of ITS sequence from each isolate was conducted against the GenBank databases.

#### Result Analysis

Genetic analysis was done using the MEGA version 5.1 software (Tamura *et al* 2006). The identity of the fungal species was established based on both the query coverage and percentage similarity of the input ITS sequence to the reference sequences in the gene bank. Reference sequence similarity of fungi above 95 % to the input ITS sequence is considered to be of same genus while similarity above 98 % is considered to be of the same species (Kumari *et al.,* 2019, Sampietro *et al* 2010). An unrooted phylogenetic tree for the DNA sequences was constructed to show relationship between fungal isolates using the neighbor joining algorithm using Clustal-W (Larkin *et al* 2007). Reference sequences for possible sequenced fungal species were included in the phylogenetic tree. Bootstrap analysis was conducted to determine the confidence for the internal nodes. Multiple sequence alignment was performed using Multalin alignment software (Corpet 1988). This alignment was performed with the aim to determine the degree of homology between DNA base sequences as well as nucleotide base composition (Whitford, 2005).

## RESULTS

### Fungal growth on plated common bean

Fungal growth was exhibited on the plates of the different stored bean cultivars after a seven days period of incubation. This was noticeable by changes in the appearance of the cultivars such as color and fruiting bodies protruding from their surfaces (Figure 3). All of the different cultivar group types were shown to be infected by fungi. Some of the plated common bean grains also had more than one fungal colony growing from them. This could be seen by differences in the colony appearance such as colors on the surfaces of the bean grains.

**Figure 3.**
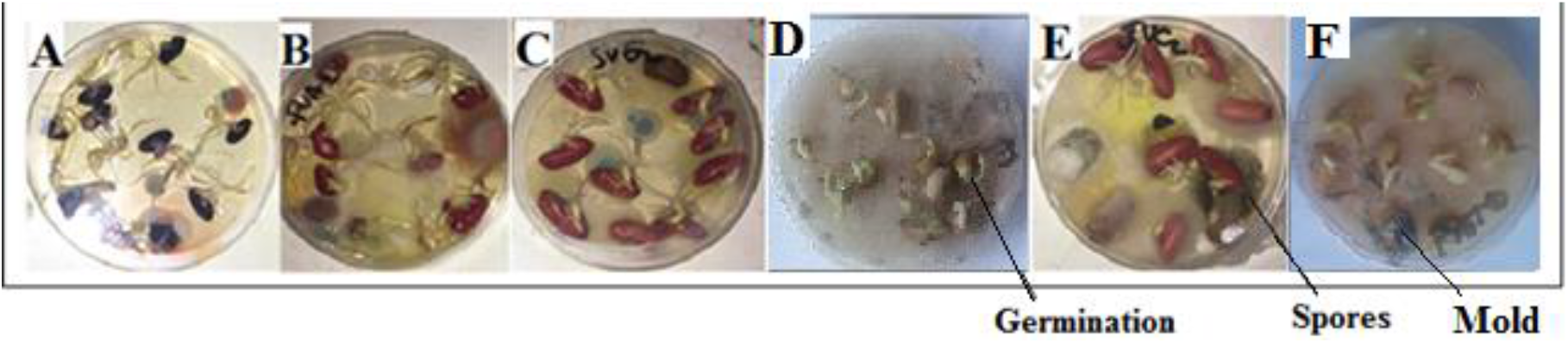
Fungal growth on plated stored bean cultivars. A- Black turtle bean (Black bean). B- Kidney bean (small sized red bean). C- Pea bean (mottled red bean). D- Navy bean (White bean) E- Large seeded bean (Large red bean). F- Pinto bean (mottled brown bean)

### Morphological characterisation fungi Isolated from stored common bean cultivars

Based on cultural characteristics (Table 1) together with micro-morphological characteristics of the fungi colonies, four distinct fungi isolate groups on the stored plated common beans were identified to be members of the genera *Aspergillus, Penicillium, Fusarium* and *‘mycelia sterilia’*.

**Table 1:**
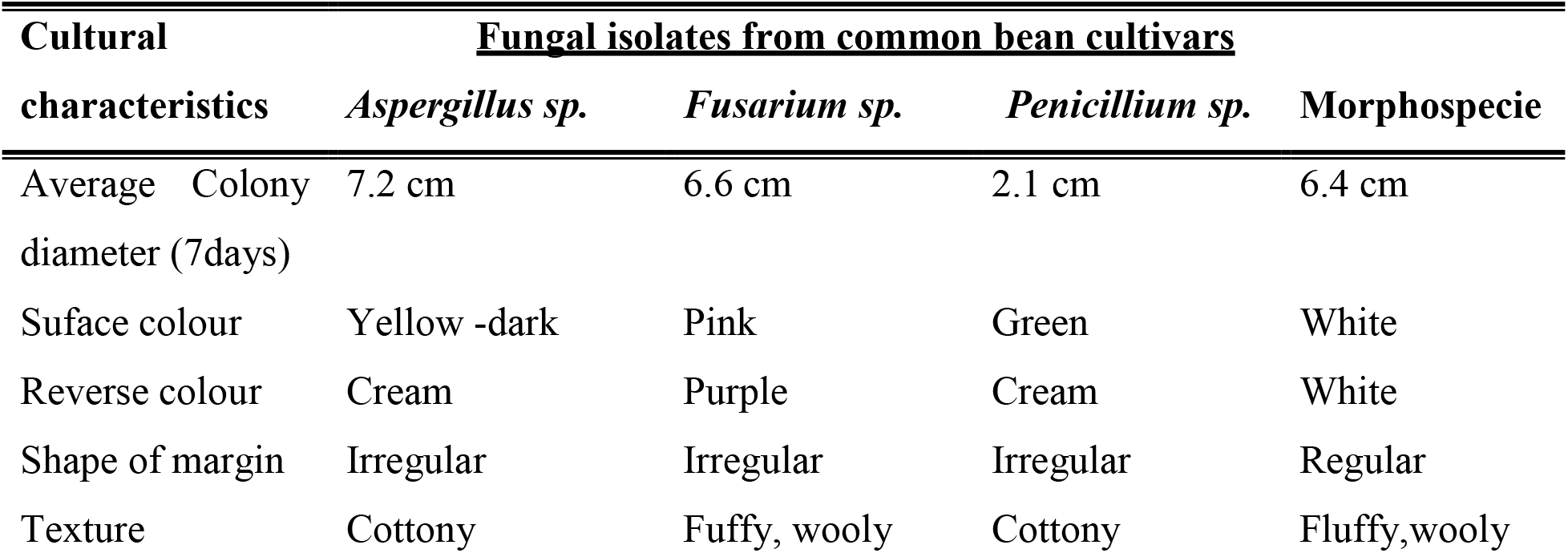
Cultural characteristics of isolated fungi on PDA media.

| Cultural characteristics | <u>Fungal isolates from common bean cultivars</u> |  |  |  |
| --- | --- | --- | --- | --- |
|  | *Aspergillus sp.* | *Fusarium sp.* | *Penicillium sp.* | Morphospecie |
| Average Colony diameter (7days) | 7.2 cm | 6.6 cm | 2.1 cm | 6.4 cm |
| Surface colour | Yellow -dark | Pink | Green | White |
| Reverse colour | Cream | Purple | Cream | White |
| Shape of margin | Irregular | Irregular | Irregular | Regular |
| Texture | Cottony | Fuffy, wooly | Cottony | Fluffy,wooly |

For *Aspergillus*, the colonies were yellow to dark on the surface and creamy on the reverse side. Texture was woolly to cottony and appeared granular. Colony was fast growing. Microcospic characteristics included long and septate cells borne on hyaline hyphae, colourless conidiophores that are wide and roughened, globose vesicles held on long conidiophores and smooth globose conidia. Conidiophores appeared dense felt with mature vesicles bearing phialides over their entire surface. (Figure 4).

**Figure 4.**
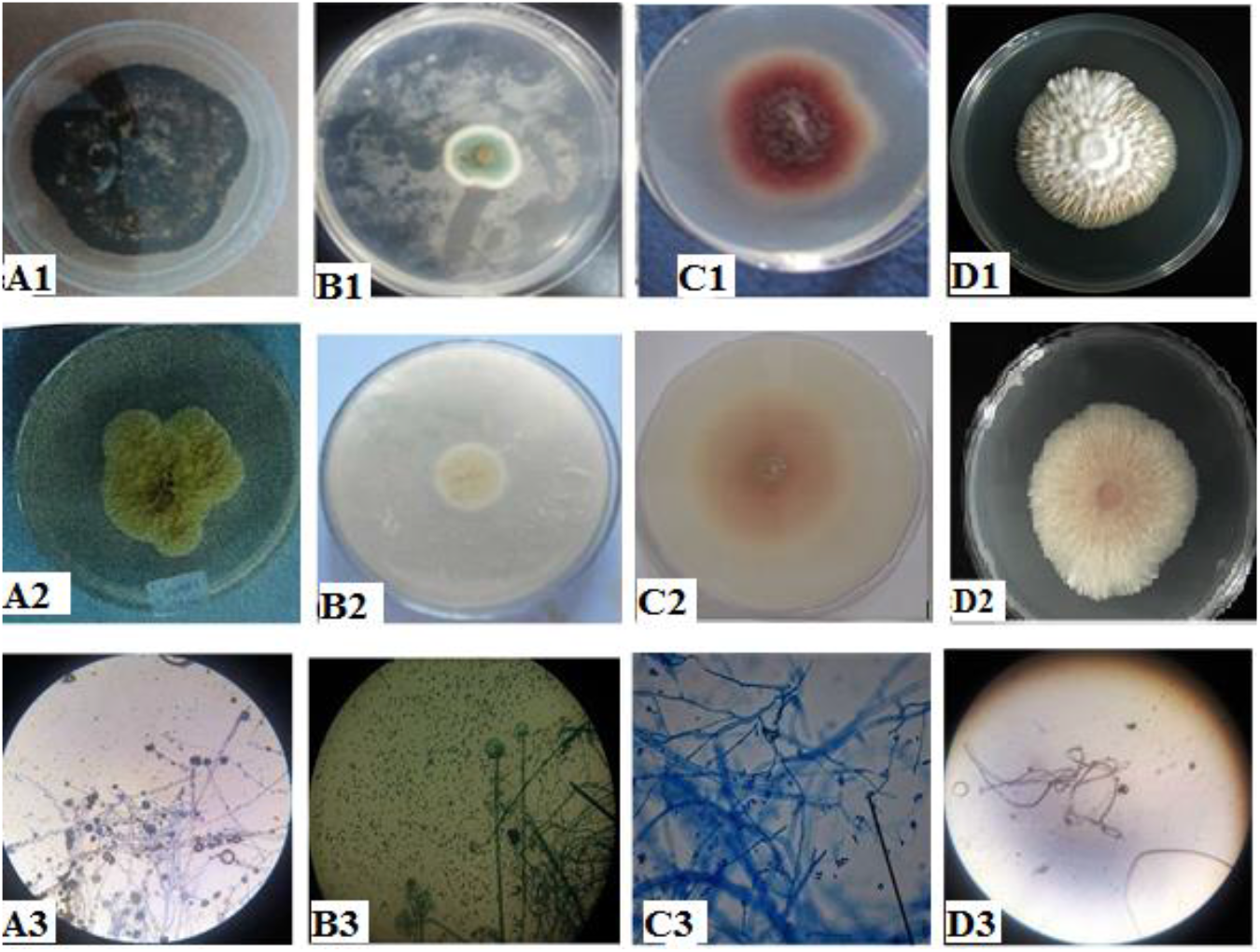
Colonial characteristics of fungal isolates (surface view A1 – D1) A1; *Aspergillus sp*. B1; *Penicillium sp*. C1; *Fusarium sp.* D1; Morphospecie **Colonial characteristics of fungal isolates (reverse view A2 – D2)** A2; *Aspergillus sp*. B2; *Penicillium sp*. C2; *Fusarium sp*. D2; Morphospecie **Microscopic images of fungi isolates A3 – D3** A3; *Aspergillus sp*. B3; *Penicillium sp*. C3; *Fusarium sp.* D3; Morphospecie

Colonies with invisible condidia/spores on stained septate mycelia on slide under the compound microscope was described as ‘Mycelia sterilia’ and termed morphospecie. Its colony appeared white on both its surface and reverse. No fruiting bodies produced (Fig. 4).

*Fusarium* colonies were fluffy, pink red on the surface and light pink on the reverse. Curved microconidia produced on simple, short phialides. Conidia had more than one cell (Figure 4).

#### Penicillium

Colonies appeared green on the surface and creamy on the reverse. Conidia were borne in unbranched chains, arising from bundles of cylindrical to bottle shaped, phialides closely arranged in a brush-like head (Figure 4)

### Molecular characterisation of fungi isolated from stored common bean cultivars

#### PCR amplification of the ITS regions

A single band was obtained after amplification of the genomic DNA, Internal Transcribed spacer (ITS) region using Universal primers (ITS1 & ITS4) for *Aspergillus, Morphospecie, Fusarium* and *Penicillium* isolates. The PCR result of the amplified ITS region in this study showed that the amplicons had an approximate size of about 600 base pairs (Figure 5).

**Figure 5:**
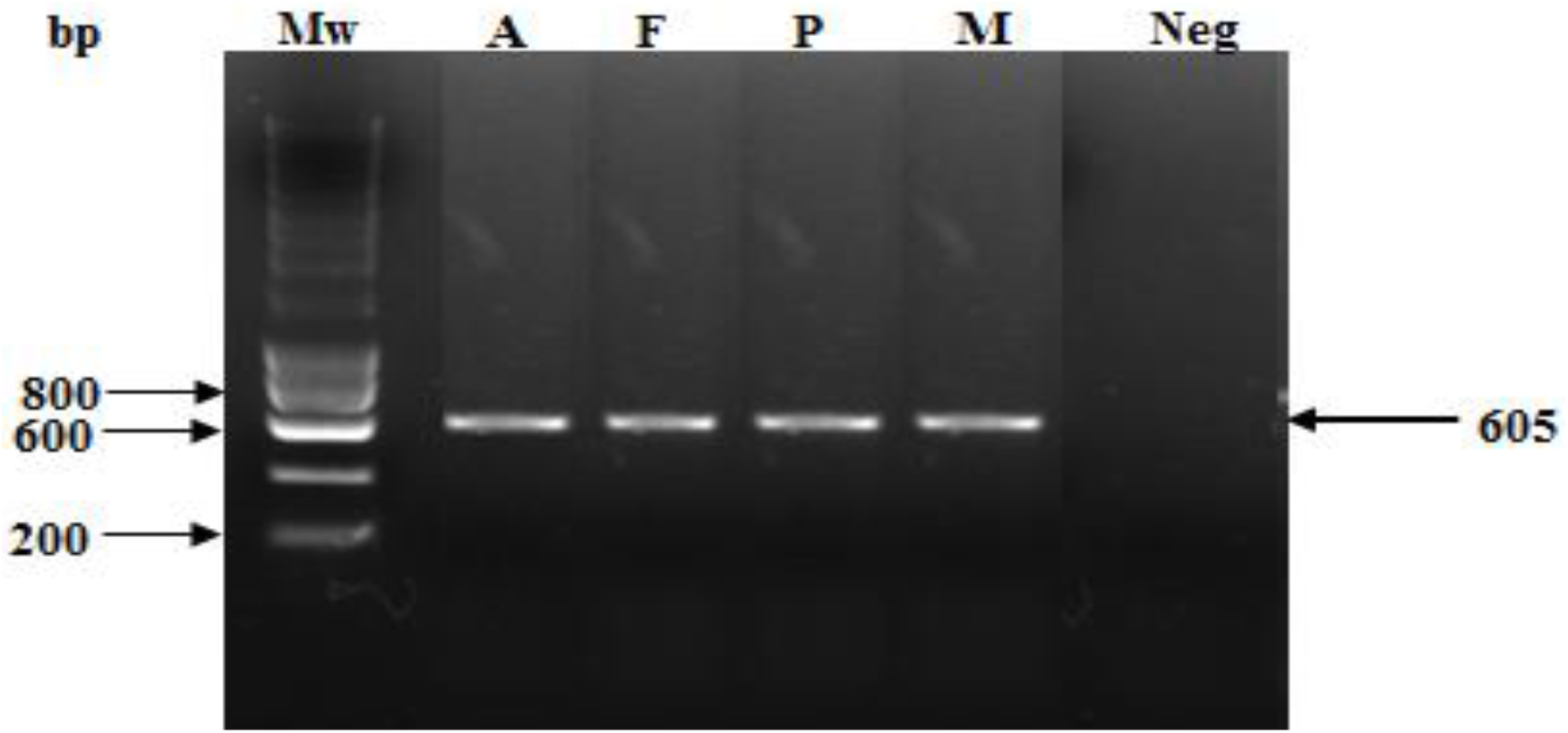
1.5% Gels showing successful amplification of fungal DNA using ITS1 and ITS4 universal primers Mw (molecular weight) = 200 bp marker A, F, P and M represent the amplicons obtained from various fungi isolate types, Neg: Negative control A= *Aspergillus sp*., F = *Fusarium sp*., P = *Penicillium sp*., M = Morphospecies

#### DNA sequencing and fungi species Identification

Identity of the fungal species were established after comparing their target ITS sequence region with reference sequences found in Genbank. Following blasting from the sequences obtained from the amplicons, the four isolates were identified as four distinct species of fungi each having E-values of zero and percentage similarity greater than 98 % to that of the gene bank reference (Table 2). These were; *Penicillium aethiopicum-* percentage identity of 99.59 % for isolate P, *Xylaria hypoxylon* - percentage identity 98.78 % for isolate M, *Aspergillus flavus* -percentage identity of 99.78 % for isolate A, *Fusarium oxysporium* - percentage identity of 99.51 % for isolate F.

**Table 2:**
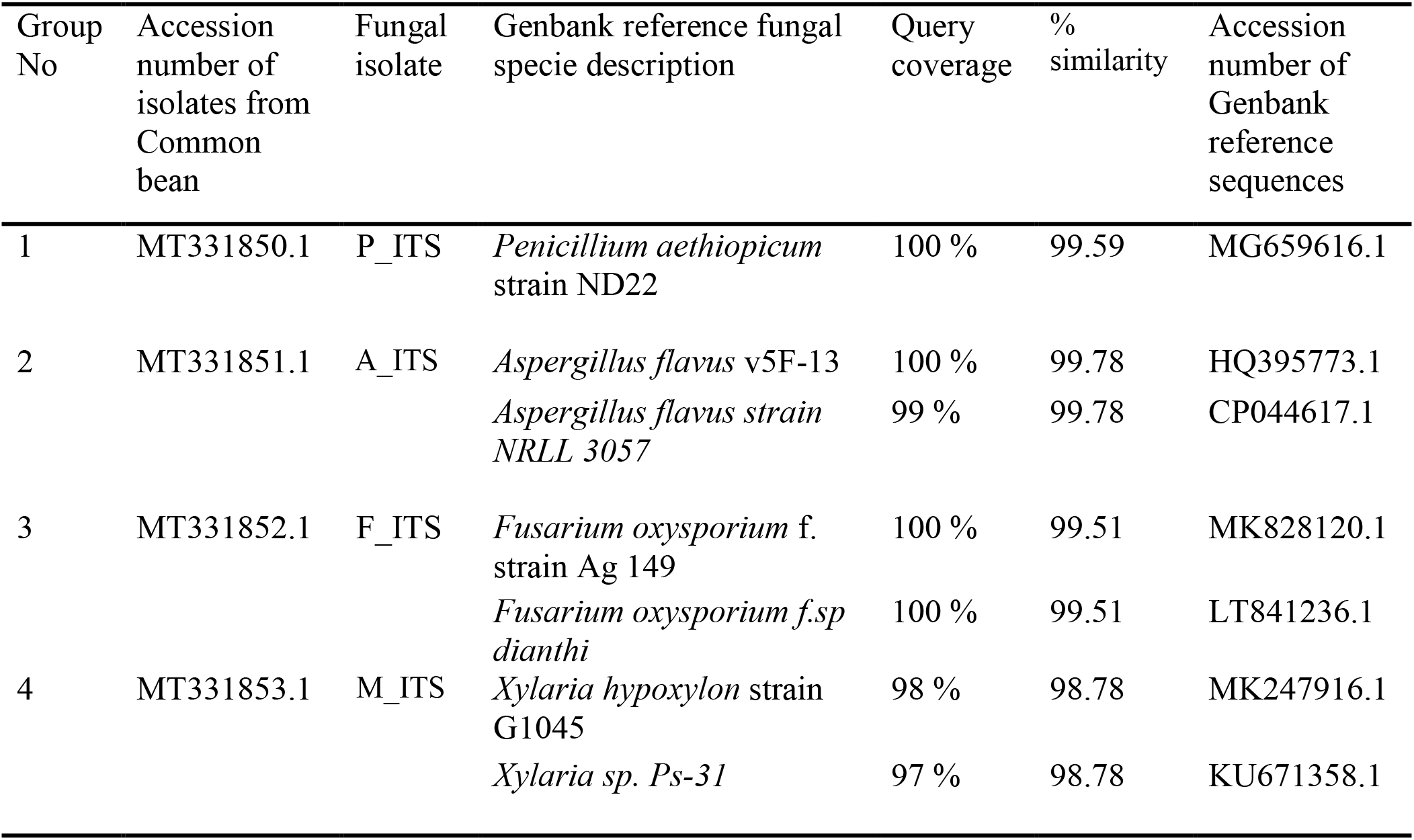
Blast results showing similarities of ITS sequences from isolates to that of species in Genbank.

| Group No | Accession number of isolates from Common bean | Fungal isolate | Genbank reference fungal specie description | Query coverage | % similarity | Accession number of Genbank reference sequences |
| --- | --- | --- | --- | --- | --- | --- |
| 1 | MT331850.1 | P_ITS | *Penicillium aethiopicum* strain ND22 | 100 % | 99.59 | MG659616.1 |
| 2 | MT331851.1 | A_ITS | *Aspergillus flavus* v5F-13 | 100 % | 99.78 | HQ395773.1 |
|  |  |  | *Aspergillus flavus* strain NRLL 3057 | 99 % | 99.78 | CP044617.1 |
| 3 | MT331852.1 | F_ITS | *Fusarium oxysporium* f. strain Ag 149 | 100 % | 99.51 | MK828120.1 |
|  |  |  | *Fusarium oxysporium* f.sp *dianthi* | 100 % | 99.51 | LT841236.1 |
| 4 | MT331853.1 | M_ITS | *Xylaria hypoxylon* strain G1045 | 98 % | 98.78 | MK247916.1 |
|  |  |  | *Xylaria* sp. *Ps-31* | 97 % | 98.78 | KU671358.1 |

The *‘mycelia sterilia’* designated morphospecie was identified based on its gene sequence similarity as *Xylaria hypoxylon*; being a member of the genus *Xylaria*, which is an ascomycetous fungi also known as sac like fungi having a typical structure called an ascus bearing 8 ascospores.

### Genetic relationship between stored fungal isolates

#### Phylogenetic relatedness of stored fungal isolates

An unrooted phylogenetic tree was produced (Figure 6) using the ITS sequence of the fungal isolates. Branch supports were compared out of 100 bootstrap replicates. Values obtained from bootstrap were quite high and ranged from 86 to 100. Bootstrap values gave the reliability of the nodes or branches. The bootstrap values due to the high value were thus robust. This showed a high confidence of support from the nodes or branches in the phylogenetic tree. The identity of the fungal isolates were shown through the same branch as the reference species obtained from the Genbank. The phylogenetic tree also showed differences between fungal species due to differences in branch lengths and nodes.

**Figure 6:**
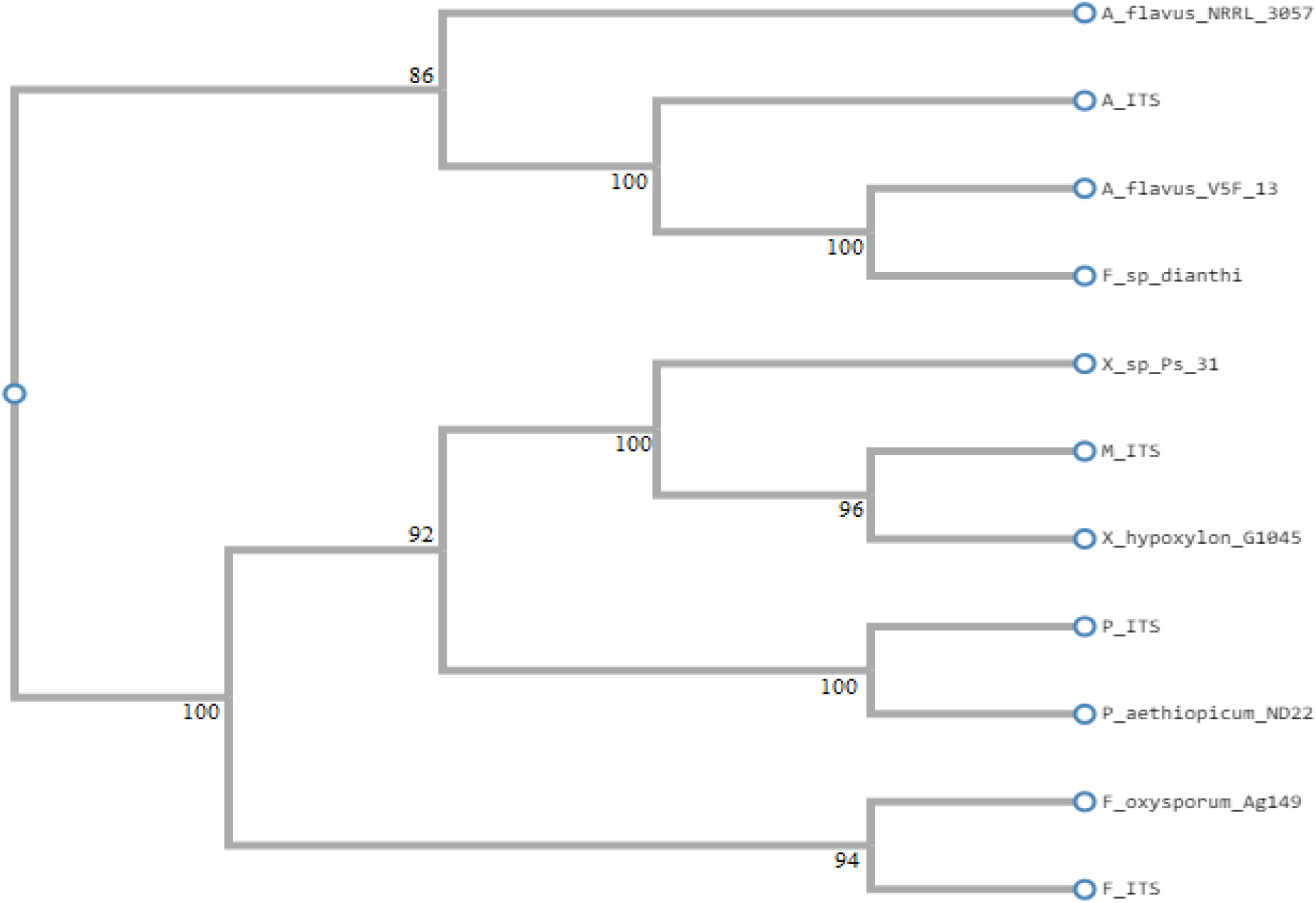
Unrooted phylogenetic tree by neighbour joining algorithm showing the relationship between fungal species obtained from stored bean cultivars. *Numbers attached to branch points shows percentage of branch supports to 100 bootstrap replicates*. Fungal isolates obtained from stored bean: M_ITS, A_ITS, F_ITS, P_ITS

#### Multiple sequence alignment analysis

To support the results of BLAST and phylogenetic analysis, the species similarity analysis of the sequence of nitrogenous bases was revealed by multi-alignment (Figure 7). The total alignment length of the sequences was 650bp.Various columns showed features of similarity and conservation. Base sequence of fungal isolates were similar to each other in sequence number 39-43, 45-49, and 261 – 390. This regions were referred to as high consensus regions along sequences column (≥ 90% alignment). Here, majority of the nucleotide residues were identical in all of the alignment sequences. The multiple alignment also featured low consensus regions (≥ 50% alignment). Region of low consensus could be highly noticed between the alignment regions 10 – 38 and 627 – 650. Gaps created between the sequence (for example; 1-10, 76 – 82, 85 – 87) represented point mismatches. At this region, the sequence identity was very small.

**Figure 7:**
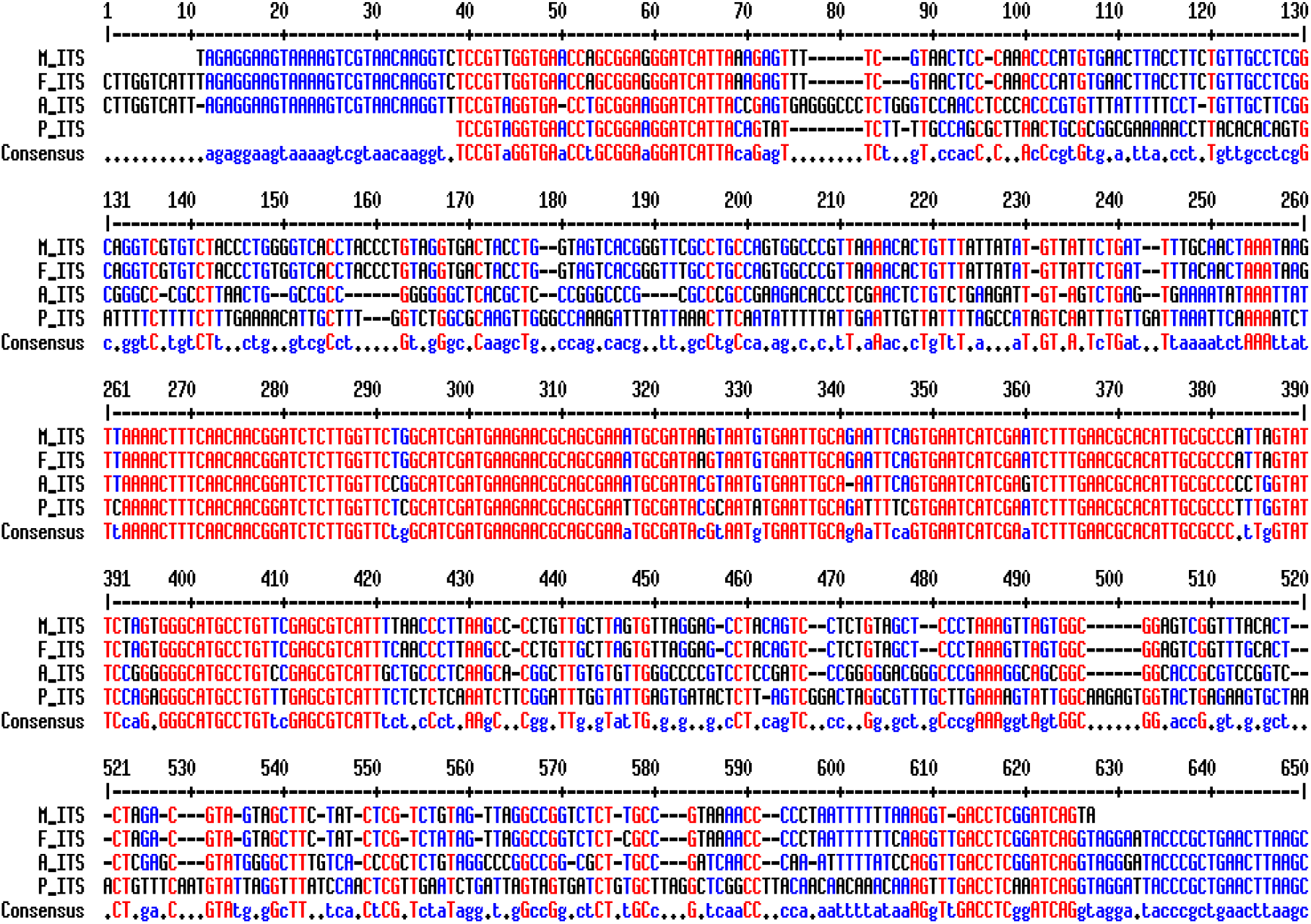
Multiple sequence alignment of fungal isolates by Multalin

## DISCUSSION

Initial characterization of fungal isolates obtained from the stored common bean cultivars using morphological techniques classified the isolates into four different genera. Molecular technique confirmed the result obtained from the morphological technique and further separated the fungal isolates into different species. The application of molecular techniques have been extensively used to identify fungi at both levels of taxonomy and phylogeny (Bruns *et al* 1991). The single band obtained in this study after DNA amplification of the ITS region of all the fungi isolates confirmed that the isolates were obtained from pure cultures. The PCR result of this amplified ITS region of the fungi was about 600 base pairs. The result of this amplification was similar to the results by (Martínez-Culebras *et al* 2009), (González-Salgado *et al* 2005) and (Patiño *et al* 2005) which were 650 bp, 612 bp, and 603 bp respectively for amplified ITS region for *Aspergillus* sp. The data resulted from BLAST demonstrated similarities between the isolated genes of the fungi obtained from the stored common bean with other species of existing fungi in Genbank. This also goes to prove successful amplification of the isolates ITS region.

Results obtained from phylogenetic analysis confirmed that the fungi recovered from the stored cultivars were all of distinct species. This possibly could be due to the nature of the conserved targeted ITS region found between 16S and 23S ribosomal RNA gene that functions in bringing about the identity of fungal species and genera (Dendis *et al* 2003). The multiple sequence alignment which was done with the objective of obtaining the areas of differences or similarities along the DNA of the isolated species showed gaps or dashes in some regions of the sequence (for example; sequence region 1-10, 76 – 82, 85 – 87). Studies indicates that the presence of these gaps in sequences is as a result of mutation either by an insertion or a deletion (Hidayat *et al* 2008).

Morphospecies are artificial groups of fungi that were believed not to reflect taxonomic relationships (Guo *et al* 2003). However, Lacap *et al* (2003) validated morphospecies as taxonomic groups based upon the analysis of the sequence of their ribosomal DNA. The identification of the morphospecies in the present research showed that they comprised of Ascomycota. The ‘*Myelia sterilia’* termed Morphospecie that couldn’t be identified based on its inability to sporulate in PDA media in which it was cultured, was however identified based on sequencing its DNA. Results produced were in close match to *Xylaria hypoxylon* (98.78 % sequence similarity). According to Rogers (2000) *Xylaria sp.* thrives well on dry food surfaces. This has been attributed to the xerophilous lifestyle of their ancestors. During storage, moisture content of stored grains decreases. This lower moisture content creates a favourable medium for the growth and survival of xerotolerant and xerophilic fungi (Iamanaka *et al* 2005).

Esiegbuya *et al* (2013) identified *Xylaria sp.* as one of the fungi species that caused postharvest dry rot disease on *Raphia hookeri* fruits. The occurence of *Xylaria* sp. has also been recorded by Parsa *et al* (2016) to be a fungal endophyte in the germinated seeds of common bean*. Xylaria hypoxylon* has also been found to be a post-harvest fungus associated with the deterioration of shea nuts and kernel (Esiegbuya *et al* 2014). There are limitations to the identification of the sterile mycelia using DNA analysis. This is because even if similarities in sequences are high between an isolate and a reference sequence, sufficient data for complete resolution are usually not always available (Promputtha *et al* 2005). Some of the isolates presented here may be new species or even genera.

DNA sequence results following BLAST analysis sequence typing also indicated that *Fusarium oxysporium* was isolated from common bean cultivars at storage in the Menoua Division, West Region of Cameroon. This was in agreement with Zhang *et al* (2006) who identified *Fusarium solani* and *Fusarium oxysporium* as two phylogenetically distinct species of *Fusarium* found in common beans. *Fusarium* requires higher moisture levels for their growth which generally occurs at harvest (Girisham *et al* 2017). The presence of *Fusarium oxysporium* in stored bean samples in Menoua Division indicated that the storage was done under poor conditions (relatively high moisture). Stored grain can be colonized by a range of different fungi that compete for space, nutrients and gas exchange and as such this antagonistic relationship limits the growth of other species. This reason could account for the significance difference (P < 0.05) in their mean occurrence on the various common bean cultivars. Marín *et al* (1998) reported that the growth of some species of *Fusarium* on stored grains were inhibited by the presence of some *Aspergillus species*.

Sequencing of DNA extracted from *Penicillium* identified *Penicillium aethiopicum* as the species found in common beans. Studies done by Njobeh *et al* (2009) showed that *Penicillium aethiopicium* occurred in all infected common bean samples isolated in Yaounde, Cameroon. *Aspergillus flavus* was also obtained from the sequenced *Aspergillus* DNA found on these stored common bean cultivars. *Aspergillus flavus* has also been reported to infect other legumes such as soya beans (Paster *et al* 1993). *Aspergillus* and *Penicillium* are widespread in nature and frequently infect grains in stores. These fungi have the tendency to survive at low levels of moisture (Pitt and Hocking 1997). Studies done by (Wrather and Sweets 2009) have described *Aspergillus* and *Penicillium* as storage fungi because they invade grains or seeds during storage

The isolation of *Aspergillus flavus* and *Fusarium oxysporum* also shows conformity with work done by Kator *et al* (2016) who isolated *Aspergillus niger, Aspergillus flavus, Fusarium oxysporum* and *Botryodiplodia theobromae* from samples of common bean obtained from selected markets in Makurdi, Nigeria. Similar studies done by Sakai *et al* (2005) shows that *Aspergillus flavus* also caused grain deterioration during storage. Differences between field and storage fungi may also play a role in determining which species will dominate a stored grain ecosystem under a particular set of environmental factors. During long-term storage, fungi of the genera *Aspergillus* and *Penicillium* (“storage flora”) progressively replace the “field flora” such as *Fusarium* and *Alternaria* over a period of several months (Fleurat *et al* 2002). *Fusarium* sp, *Aspergillus* sp and *Penicillium* sp has also been shown to be associated with cola nuts, groundnuts and sweet potato in the western highlands of Cameroon (Ngoko *et al* 2008).

## CONCLUSION

In this study, fungi isolated from stored common bean grains was identified based on their phenotypic features (Morphological characterisation) and genotypic features (Molecular characterisation). These species isolated are a significant proof that contamination of common beans occurs during storage. This study therefore brings forth a novel information which will assist in establishing measures to prevent fungal infection of stored common bean in the Menoua Division.

## ACKNOWLEDGEMENTS

The authors are thankful for the support provided by farmers of Menoua Division for the provision of common bean cultivars.

## REFERENCES

1. Bechem, E. T. and Afanga, Y. A. (2017). Morphological and molecular identification of fungi associated with corm rot and blight symptoms on plantain (Musa paradisiaca) in macro-propagators. International Journal of Biological and Chemical Sciences, 11(6), 2793–2808.

2. Beebe, S. E., Rao, I. M., Mukankusi, C. M. and Buruchara, R. A. (2012). Improving resource use efficiency and reducing risk of common bean production in Africa, Latin America, and the Caribbean. Centro Internacional de Agricultura Tropical (CIAT).

3. Bruns, T. D., White, T. J., and Taylor, J. W. (1991). Fungal molecular systematics. Annual Review of Ecology and systematics, 22(1), 525–564.

4. Corpet, F. (1988). Multiple sequence alignment with hierarchical clustering. Nucleic acids research, 16(22), 10881–10890.

5. Dendis, M., Horvath, R., Michalek, J., Růžička, F., Grijalva, M., Bartoš, M. and Benedík, J. (2003). PCR-RFLP detection and species identification of fungal pathogens in patients with febrile neutropenia. Clinical Microbiology and Infection, 9(12), 1191–1202.

6. Esiegbuya, D.O., Okungbowa, F.I., Oruade-Dimaro, E.A and Airede, C.E. (2013). Dry rot of Raphia hookeri fruits and its effect on the mineral and proximate composition. Nigerian Journal of Biotechnology, 26 (4), 26–32.

7. Esiegbuya, D., Osagie, J., Okungbowa, F. and Ekhorutomwen, E. (2014). Fungi Associated with the Postharvest Fungal Deterioration of Shea Nuts and Kernels. International Journal of Agriculture and Forestry 4 (5), 373–379.

8. Fleurat-Lessard, F. (2002). Qualitative reasoning and integrated management of the quality of stored grain: a promising new approach. Journal of Stored Products Research, 38(3), 191–218.

9. Ferrer, C., Colom, F., Frasés, S., Mulet, E., Abad, J. L. and Alió, J. L. (2001). Detection and identification of fungal pathogens by PCR and by ITS2 and 5.8 S ribosomal DNA typing in ocular infections. Journal of clinical microbiology, 39(8), 2873–2879.

10. Girisham, S., Rao, V. K. and Reddy, S. (2017). Characterisation of pathogenic fungi on grains. Taxonomy of mycotoxigenic fungi, 2(3), 86–91 Scientific Publishers.

11. González-Salgado, A., Patiño, B., Vázquez, C. and González-Jaén, M. T. (2005). Discrimination of *Aspergillus niger* and other *Aspergillus* species belonging to section Nigri by PCR assays. FEMS Microbiology Letters, 245(2), 353–361.

12. Guo, L., Hyde, K. and Liew, E. (2000). Identification of endophytic fungi from Livistona chinensis based on morphology and rDNA sequences. The New Phytologist, 147(3), 617–630.

13. Hidayat, T., Kusumawaty, D., Kusdianti, Y. D., Muchtar, A.A. and Mariana, D. (2008). Using the base sequence of DNA Internal Transcribed Spacer Region (ITS). Education Department of Biology, Faculty of Mathematics and Science Education University of Indonesia, Bandung

14. Iamanaka, B. T., Taniwaki, M. H., Menezes, H. C., Vicente, E. and Fungaro, M. H. P. (2005). Incidence of toxigenic fungi and ochratoxin A in dried fruits sold in Brazil. Food additives and contaminants, 22(12), 1258–1263.

15. Kamtchoum, S. M., Nuemsi, P. P. K., Tonfack, L. B., Edinguele, D. G. M. and Kouahou, W. N. (2018). Production of Bean (*Phaseolus vulgaris L*.) under organo-mineral fertilization in humid forest agro-ecological zone with bimodal rainfall pattern in Cameroon. Annual Research and Review in Biology, 4(2) 1–11

16. Kator, L., Aondo, T. and Akinyemi, B. (2016). Isolation and identification of seed borne fungi of common Bean (*Phaseolus vulgaris* L.) from selected markets in Makurdi. International Journal of Applied Agricultural Sciences, 2(5), 75–78.

17. Klich, M. A. (2002). Identification of common Aspergillus species.

18. Kumar, D. and Kalita, P. (2017). Reducing Postharvest Losses during Storage of Grain Crops to Strengthen Food Security in Developing Countries. Food Journal, 6(1), 15 – 22.

19. Kumari, R., Jayachandran, L. E. and Ghosh, A. K. (2019). Investigation of diversity and dominance of fungal biota in stored wheat grains from governmental warehouses in West Bengal, India. Journal of the Science of Food and Agriculture, 99(7), 3490–3500.

20. Lacap, D., Hyde, K. and Liew, E. (2003). An evaluation of the fungal ‘morphotype’ concept based on ribosomal DNA sequences. Fungal Diversity.

21. Larkin, M. A., Blackshields, G., Brown, N., Chenna, R., P. A. and McWilliam, H. (2007). Clustal W and Clustal X version 2.0. Bioinformatics, 23(21), 2947–2948.

22. Marin, S., Sanchis, V., Ramos, A. J., Vinas, I. and Magan, N. (1998). Environmental factors, in vitro interactions, and niche overlap between *Fusarium moniliforme*, *F. proliferatum*, and *F. graminearum*, *Aspergillus* and *Penicillium* species from maize grain. Mycological Research, 102(7), 831–837.

23. Martínez-Culebras, P., Crespo-Sempere, A., Sánchez-Hervás, M., Elizaquivel, P. and Aznar, R. (2009). Molecular characterization of the black Aspergillus isolates responsible for ochratoxin A contamination in grapes and wine in relation to taxonomy of *Aspergillus* section Nigri. International journal of food microbiology, 132(1), 33–41.

24. Nega, A. (2014). Isolation and identification of fungal pathogens associated with cold storage type of (Coffee arabica) seed, at Jimma agricultural research center, Western Ethiopia. Journal of Biology and Agricultural Healthcare, 4 (2), 20–26.

25. Njobeh, P. B., Dutton, M. F., Koch, S. H., Chuturgoon, A., Stoev, S. and Seifert, K. (2009). Contamination with storage fungi of human food from Cameroon. International Journal of Food Microbiology, 135(3), 193–198.

26. Ngoko, D.H., Imele, P.T and Kamga1, S. M. (2008). Fungi and mycotoxins associated with food commodities in Cameroon. Journal of Applied Biosciences, 6(2): 164 – 168

27. Omotayo, O. P., Omotayo, A. O., Mwanza, M. and Babalola, O. O. (2019). Prevalence of mycotoxins and their consequences on human health. Toxicological research, 35(1), 1–7.

28. Parsa, S., García-Lemos, A. M., Castillo, K., Ortiz, V. and López-Lavalle, L. A. (2016). Fungal endophytes in germinated seeds of the common bean, *Phaseolus vulgaris*. Fungal biology, 120(5), 783–790.

29. Paster, N., Droby, S., Chalutz, E., Menasherov, M., Nitzan, R. and Wilson, C. (1993). Evaluation of the potential of the yeast Pichia guilliermondii as a biocontrol agent against Aspergillus flavus and fungi of stored soya beans. Mycological Research, 97(10), 1201–1206.

30. Patiño, B., González-Salgado, A., González-Jaén, M. T. and Vázquez, C. (2005). PCR detection assays for the ochratoxin-producing *Aspergillus carbonarius* and *Aspergillus ochraceus* species. International Journal of Food Microbiology, 104(2), 207–214.

31. Pitt, J. and Hocking, A. D. (2009). Methods for Isolation, Enumeration and Identification in n Fungi and Food Spoilage (19–52). Springer, Boston, MA.

32. Promputtha, I., Jeewon, R., Lumyong, S., McKenzie, E. H. and Hyde, K. D. (2005). Ribosomal DNA fingerprinting in the identification of non sporulating endophytes from *Magnolia liliifera* (Magnoliaceae). Fungal Diversity, 18(24), 61–66

33. Richard, J. L. (2007). Some major mycotoxins and their mycotoxicoses—An overview. International journal of food microbiology, 119(1-2), 3–10.

34. Rogers, J. D. (2000). Thoughts and musings on tropical Xylariaceae. Mycological Research, 104(12), 1412–1420.

35. Sakai, A., Tanaka, H., Konishi, Y., Hanazawa, R. and Ota, T. (2005). Mycological examination of domestic unpolished rice and mycotoxin production by isolated *Penicillium islandicum*. Shokuhin eiseigaku zasshi. Journal of the Food Hygienic Society of Japan, 46(5), 205–212.

36. Sampietro, D.A., Marin, P., Iglesias, J., Presello, D.A. and Vattuone, M.A. (2010). A Molecular based strategy for rapid diagnosis of toxigenic *Fusarium* species associated to cereal grains from Argentina. Fungal Biology. 114(4), 74–81.

37. Shende, K. S. and Lifeter, Y. B. (2017). Post-harvest challenges of food crops in Jakairi subdivision, Cameroon. A threat to food security. International Journal of Agriculture and Environmental Research, 7(10), 2455–6939.

38. Tamura, K., Dudley, J., Nei, M. and Kumar, S. (2006). Molecular Evolutionary Genetics Analysis (MEGA).

39. Whitford, D. (2005). Protein : Structure and Function. John Wiley and Sons Ltd. England.

40. Wrather, J. A. and Sweets, L. (2009). Management of Grain Sorghum Diseases in Missouri. Extension publications (MU).

41. Zhang, N., O’Donnell, K., Sutton, D. A., Nalim, F. A. and Summerbell, R. C. (2006). Members of the *Fusarium solani* species complex that cause infections in both humans and plants are common in the environment. Journal of Clinical Microbiology, 44(6), 2186–2190.

42. Zhang, Y., Zhang, S., Liu, X., Wen, H. and Wang, M. (2010). A simple method of genomic DNA extraction suitable for analysis of bulk fungal strains. Letters in applied microbiology, 51(1), 114–118.

